# eIF2Bε phosphorylation activates integrated stress response-mediated protein translation and links stress signalling to amyloidogenesis in Alzheimer’s disease

**DOI:** 10.1101/2025.11.03.686297

**Authors:** Hugo Fanlo-Ucar, Clara Planas-Clopes, Oriol Bagudanch, Carlos Pardo-Pastor, Scarlett Troncoso-Cotal, Alejandra R. Alvarez, Baldo Oliva, Francisco J. Muñoz

## Abstract

A central etiopathogenic event in Alzheimer’s disease (AD) is the accumulation of amyloid β-peptide (Aβ) derived from the amyloidogenic processing of the amyloid precursor protein, a pathway initiated by BACE1. Chronic activation of the Integrated Stress Response is linked to AD, typically through eIF2α phosphorylation, which selectively enhances translation of key stress-responsive mRNAs like ATF4 and BACE1. We investigated a novel regulatory mechanism mediated by Glycogen Synthase Kinase-3β (GSK-3β) a hyperactive kinase in AD, on the translational factor eIF2B, the guanine nucleotide exchange factor (GEF) for eIF2. The methodology combined cellular and molecular biology approaches (western blot, ELISA, immunofluorescence, and luciferase assays) using pharmacological inhibitors and plasmid transfection in established cell models, along with computational structural modeling (AlphaFold3). Findings were validated in post-mortem human brain tissue, with statistical analyses (t-tests and ANOVA) applied throughout. We treated cells (SH-SY5Y) with a GSK-3β inhibitor (NP031112) or overexpressed a constitutively active GSK-3β mutant (GSK-3β-S9A) to modulate GSK-3β activity. GSK-3β inhibition significantly reduced ATF4 and BACE1 protein levels in SH-SY5Y cells in a dose-dependent manner as we tested by western blot (*P*<0.001), independent of eIF2α phosphorylation. This effect was translational, as the inhibitor still reduced BACE1 levels when transcription was blocked. Crucially, NP031112 reversed stress-induced eIF2Bε phosphorylation at Ser540 (*P*<0.001). Furthermore, H_2_O_2_-induced Aβ_1-42_ secretion by SH-SY5Y cells was significantly reversed by GSK-3β inhibition (*P*<0.001). Luciferase reporter assays using the BACE1 5’ untranslated region (5’ UTR) confirmed that both eIF2Bε silencing and a phosphomimetic mutant eIF2Bε-S540E increased BACE1 5’ UTR-driven translation (*P*<0.001), demonstrating that reduced functional eIF2B enhances BACE1 translation. Finally, immunohistofluorescence and western blot analysis of human AD hippocampi showed a significant increase in pS540-eIF2Bε, ATF4, and BACE1 levels in AD patients compared to non-demented controls (*P*<0.001 for all three). Our findings establish a novel GSK-3β-eIF2Bε-BACE1 axis that links stress response dysregulation to amyloidogenesis, independent of canonical eIF2α phosphorylation. The phosphorylation of eIF2Bε functionally mimics the effects of eIF2α phosphorylation by reducing the available functional eIF2B pool, thereby reprogramming translation to favour the production of stress proteins like ATF4 and BACE1. We proposed eIF2Bε phosphorylation is a key etiological mechanism in AD.

## Introduction

A hallmark of Alzheimer’s disease is the accumulation of misfolded amyloid β-peptide (Aβ) peptides, produced through amyloidogenic processing of the amyloid precursor protein (APP). In this pathway, BACE1 initiates APP cleavage,^1^ followed by γ-secretase,^2,3^ which determines Aβ length. These peptides aggregate into toxic oligomers and fibrils, impairing synaptic function and neuronal viability.^4,5^ However, how amyloidogenic processing becomes activated with age, the major Alzheimer’s disease risk factor, remains unclear.

In Alzheimer’s disease and other neurodegenerative diseases, chronic integrated stress response (ISR) activation disrupts proteostasis, synaptic function and survival.^6–8^ Dysregulation of ISR components, particularly eIF2α phosphorylation and ATF4-mediated transcription, is linked to protein misfolding,^9,10^ oxidative stress,^11^ and neuroinflammation.^12^ ATF4 regulates genes controlling adaptation, metabolism and apoptosis,^13^ but under prolonged stress it promotes neuronal death.^14^ The key regulatory event is eIF2α phosphorylation, which reduces general translation but enhances synthesis of uORF-containing mRNAs,^15^ such as ATF4 and BACE1.^16–20^ By lowering ternary complex availability, phosphorylated eIF2α slows scanning and reinitiation, increasing translation of ATF4 and BACE1, the latter contributing to Aβ overproduction.^13,17^

eIF2B, the GEF of eIF2, is central in this process.^21^ Normally, it replenishes eIF2-GTP to sustain translation. Under stress, phosphorylated eIF2α binds eIF2B with high affinity, sequestering it in an inactive state and reducing protein synthesis.^13^ This allows global suppression while maintaining selective translation of stress-response proteins. eIF2B activity is further regulated by GSK-3β,^22,23^ a kinase hyperactive in Alzheimer’s disease due to Aβ accumulation.^24,25^ GSK-3β promotes tau hyperphosphorylation, neuronal dysfunction and possibly ISR modulation. It phosphorylates the ε subunit of eIF2B at Ser540, reducing GEF activity by ∼50%.^22,23^ Although GSK-3β has been linked to BACE1 transcriptional regulation via NF-κB,^26,27^ its role in ISR-driven translational control of ATF4 and BACE1 remains unknown.

In this work we hypothesized that eIF2Bε phosphorylation significantly decreases active eIF2B, enhancing ATF4 and BACE1 expression under stress. We aim to unveil a novel translational regulatory mechanism contributing to neurodegeneration in Alzheimer’s disease.

## Materials and methods

### Reagents

GSK-3β inhibitor (NP031112) was purchased from SelleckChem (S2823). DRB was purchased from Sigma (D1916-50MG). Hydrogen peroxide solution was purchased from Sigma (216763-500ML-M). MTT (3-(4,5-dimethylthiazol-2-yl)-2,5-diphenyltetrazolium bromide) was purchased from Sigma (M5655).

### Cell lines

Human neuroblastoma cells (SH-SY5Y) were grown with Ham’s F12 medium + GlutaMAX (Gibco) supplemented with 15% FBS (Biowest) and 1% penicillin/streptomycin (Gibco). Human embryonic kidney 293 cells (HEK-293) were grown with DMEM medium (Gibco) supplemented with 10% foetal bovine serum and 1% penicillin/streptomycin. Cells were incubated at 37°C in a humidified atmosphere of 5% CO_2_.

SH-SY5Y were plated on 96-well plates (15.000 cells/well) for MTT viability assays, on 6-well plates (400.000 cells/well) for WB studies and on 90 mm Petri dishes (1.500.000 cells/plate) for the ELISA assay. HEK-293 cells were seeded 96-well plates (16.000 cells/well) for luciferase assays and in 6-well plates (300.000 cells/well) for transfection and WB studies.

### Human hippocampal samples

Human hippocampal samples were supplied by the Neurological Tissue Bank of the Biobank Hospital Clínic-IDIBAPS, Barcelona, Spain. The procedure was carried out according to the rules of the Helsinki Declaration and to the Ethics Committee of the Institut Municipal d’Investigacions Mèdiques-Universitat Pompeu Fabra (EC-IMIM-UPF). Samples were obtained from six non-demented controls (3 women, 3 men; mean age 70.2 ± 4.3 years) and six Alzheimer’s disease patients (3 women, 3 men; mean age 72.0 ± 1.3 years).

### Ethics and Consent to Participate

Human hippocampal samples were supplied by the Neurological Tissue Bank of the Biobank-Hospital Clínic-IDIBAPS (Barcelona, Spain). The Neurological Tissue Bank obtained all samples post-mortem and under written informed consent from the participants’ legal next-of-kin, in compliance with the relevant EU and national ethical guidelines and Hospital Clínic regulations.

The samples were provided to us in a fully anonymized manner. Only relevant anonymized clinical data (sex, age at death, and associated pathology/pathologies) were provided alongside the samples, strictly following the Brain Bank’s regulations and ensuring the protection of the donors’ privacy in accordance with the ethical standards observed.

In our laboratory at UPF, the research protocol was reviewed and carried out in accordance with the principles of the Declaration of Helsinki and the regulations of the Ethics Committee of the Institut Municipal d’Investigacions Mèdiques-Universitat Pompeu Fabra (EC-IMIM-UPF). Specifically, the use of these samples for this research protocol was reviewed and approved by the EC-IMIM-UPF with the reference number 2022/10334/I.

### MTT viability assay

SH-SY5Y cells seeded in a 96-well plate were treated with vehicle (DMSO) and GSK-3β inhibitor at rising concentrations both in FBS-free Ham’s F12 medium for 24 h. Then, cells were incubated with 0.5 mg/mL MTT solution for 2 h. Formazan crystals were solubilized in 100 µL DMSO and cell viability was determined by absorbance at 595 nm by VICTOR Nivo Multimade Plate Reader (PerkinElmer).

### Immunoblotting

SH-SY5Y cells in 6-well plates kept/treated with FBS-containing Ham’s F12, FBS-free Ham’s F12 or drugs (in FBS-free Ham’s F12) for 24 h; HEK-293 cells in 6-well plates transfected with 2.5 μg of GSK-3β-S9A/GSK-3β-WT constructs for 24 h; and hippocampal samples from control and Alzheimer’s disease patients, were lysed on ice with 50 µL of lysis buffer (150 mM Sodium Chloride, 1.0% Triton X-100, 0.5% Sodium Deoxycholate, 0.1% sodium dodecyl sulphate (SDS), 50 mM Tris, pH 8.0) supplemented with phosphatase and protease inhibitors tablets. Samples were centrifuged at 15,000 g for 10 min at 4°C and then loaded into 10% acrylamide protein gels. Next, proteins were transferred onto 0.2 µm pore nitrocellulose membranes. Membranes were blocked for 1 h at room temperature (RT) with either 5% bovine serum albumin (BSA) for phosphoproteins or 5% skimmed milk for total proteins in Tween 20-Tris buffer solution (TTBS: 100 mM Tris-HCl, 150 mM NaCl, pH 7.5 plus 0.1% Tween-20). Then, membranes were incubated overnight (o.n.) at 4°C with anti-BACE1 (1:1,000; ThermoFisher, PA1-757), anti-ATF4 (1:1,000; Cell Signaling, 11815S), anti-pS9-GSK-3β (1:1,000; Cell Signaling, 9323S), anti-GSK-3β (1:1,000; Cell Signaling, 9832), anti-pS51-eIF2α (1:1,000; Cell Signaling, 3597S), anti-eIF2α (1:1,000; Cell Signaling, 2103), anti-pS540-eIF2Bε (1:1,000; Invitrogen, 44-530G), anti-eIF2Bε (1:1,000; Cell Signaling, 3595S) or anti-GAPDH (1:2,000; Abcam, ab8245) antibodies (Abs). Horseradish peroxidase-conjugated donkey anti-rabbit and anti-mouse Abs (GE Healthcare) were used as secondary Abs at 1:2,000 for 1 h at RT. Primary and secondary Abs were diluted in blocking solution. Bands were visualized with Clarity Western ECL Substrate (BioRad) and analysed with ImageJ software.

### Human Aβ_1-42_ ELISA assay

Approximately 5 mL of supernatants were collected from SH-SY5Y cells cultured in 90 mm dishes and treated with vehicle, 10 μM NP031112, 10 μM H_2_O_2_ or both. These samples were centrifuged at 1200 rpm for 5 min to remove any floating cells, and concentrated 60x using the AMICON 3k filters (Sigma). Assay was performed with the commercial Ultrasensitive Amyloid beta 42 Human ELISA Kit from Invitrogen following their protocol. Final Aβ concentrations were relativized to total protein concentration.

### Structural modelling of eIF2-eIF2B

Structural models of eIF2 in complex with selected eIF2B subunits were generated using AlphaFold3. Each prediction included the eIF2 subunits α, β and γ, and the γ and ε subunits of eIF2B; the remaining eIF2B subunits were excluded from the modelling. The complete eIF2B structure was reconstructed by superimposing the modelled components onto the eIF2B structure from PDB entry 6K71. A GTP molecule was included in AlphaFold3 modelling which position it in the nucleotide-binding pocket of eIF2γ. To mimic regulatory phosphorylation, serine residues S525, S535 and S540 in eIF2Bε were substituted with glutamate prior to model generation (thus, two models were obtained, one phosphorylated and another unphosphorylated).

To assess the potential interaction between tRNA and the modelled eIF2-eIF2B complex, a tRNA molecule was incorporated by aligning the modelled eIF2 structure with the eIF2-tRNA complex from PDB entry 3JAP using the Matchmaker tool in ChimeraX. Conformational states of eIF2β and its interactions with GTP, the HEAT domain of eIF2Bε, and tRNA were analysed in ChimeraX. Electrostatic interactions were inferred based on the spatial proximity of acidic and basic side chains.

### Cloning of BACE1 5′-untranslated region

Total RNA was extracted from SH-SY5Y cells, and one-step RT-PCR was carried out using a One-Step PCR kit (Qiagen) with primers designed to amplify *BACE1* 5′ UTR: FW 5′-TGGTACCACAAGTCTTTCCGCCTCCCCA-3′, RV 5′-GAAGCTTGGTGGG CCCCGGCCTTC-3′. The PCR product, a single band matching the molecular weight of *BACE1* 5′ UTR (∼460 nt), was isolated and purified from an agarose gel using the Illustra™ GFX™ PCR DNA and Gel Band Purification kit (GE Healthcare). The 5′ UTR DNA fragment was then inserted into a modified pGL4.10-*luc2* vector from Promega containing the SV40 promoter, between Kpn1 and HindIII restriction enzyme sites.

### Transient DNA transfection of HEK-293 cells

For the overexpression studies, HEK-293 cells seeded in 6-well plates and transfected with 2.5 μg of either pcDNA3 or GSK-3β-S9A/GSK-3β-WT constructs for 24 h using the Lipofectamine 3000 transfection reagent (ThermoFisher). GSK-3β constructs were gently provided by Félix Hernández Pérez (Universidad Autónoma de Madrid, Madrid).

For the luciferase assays, HEK-293 cells were seeded in 96-well plates at a density of 12,000 cells per well and grown for 24 h with DMEM plus 10% FBS and at the same time transfected with either siCtrl or sieIF2Bε (100 pmol) using lipofectamine RNAiMAX transfection reagent (ThermoFisher). After 24 h, a total of 100 ng of DNA were transfected into each well, adjusting to the following conditions: 33 ng of either pGL4.10-*luc2* or pGL4.10-5’ UTR·*BACE1*, plus 33 ng of pcDNA3, eIF2Bε-WT or eIF2Bε-S540E, plus 33 ng of Renilla luciferase. Cells were transfected using Lipofectamine 3000 transfection reagent (ThermoFisher) for 24 h.

### Luciferase assay

Luciferase and Renilla activities of previously transfected HEK-293 cells were measured by using the Dual-Glo™ Luciferase Assay System (Promega) following the manufacturer instructions, and luminescence was read using a luminometer (VICTOR Nivo, PerkinElmer).

### Immunohistofluorescence assays

Human hippocampal sections were treated with 0.15 M glycine and 10 mg/mL NaBH_4_ in PBS for 30 min. Then they were washed in deionized water, permeabilized with 0.3% Triton X-100 in PBS for 1 h at 4°C, and then blocked with 5% FBS + 1% BSA diluted in 0.1% Triton X-100 in PBS for 2 h. The following step was incubation o.n. at 4 °C with anti-pS9-GSK-3β (1:200; Cell Signaling, 9323S) or anti-pY216-GSK-3β (1:200; Abcam, ab75745), and anti-GSK-3β (1:250; Cell Signaling, 9832) Abs. Samples were incubated the next day with Alexa Fluor 488 donkey anti-rabbit Ab (1:2000; Invitrogen, A32790), Alexa Fluor 555 donkey anti-mouse Ab (1:2000; Invitrogen, A32773) and TO-PRO-3 (1:1000) for 1 h at RT. They were then treated for 10 min with 0.3% Sudan Black (Sigma) in ethanol that was previously mixed o.n. at 4°C, centrifuged at 3000 g for 20 min and filtered using common filter paper. Sections were washed with deionized water and cover slips were mounted using Fluoromont G. Digital images were taken with a Leica TCS SP5 confocal microscope and analysed with ImageJ software.

### Statistical analyses

Data are expressed as mean ± SD of n experiments, as indicated in the corresponding figures. All statistics were performed with GraphPad software (9.0.1). Statistical tests to assure normality of the data were performed first, including Anderson-Darling, D’Agostino-Pearson omnibus, Shapiro-Wilk and Kolmogorov-Smirnov. Then, statistical analyses were performed using two-tailed Student’s t-test and one-way or two-way ANOVA, depending on the variables analysed, followed by either Dunnett’s or Tukey’s post hoc test, as appropriate for the specific experimental design and comparisons required.

## Results

### The ISR up-regulates ATF4 and BACE1 protein levels via GSK-3β activation

Serum deprivation is known to activate the ISR.^28^ To evaluate this effect in SH-SY5Y cells, we examined the expression levels of ATF4, since it is the major ISR downstream effector, and BACE1, a key enzyme in Alzheimer’s disease aetiology, after 24 h of serum starvation. A significant increase in the expression of both proteins was observed under serum-free conditions (*P* < 0.001) (Fig. 1A). Furthermore, GSK-3β, another key enzyme in Alzheimer’s disease etiopathology, is tightly regulated by phosphorylation: it becomes enzymatically enhanced via autophosphorylation at Tyr216,^29,30^ and is inhibited by phosphorylation at Ser9 by mainly Akt.^31^ Interestingly, it has been reported that serum deprivation activates GSK-3β in cortical neurons ^32^. Therefore, we investigated the role of GSK-3β in the upregulation of ATF4 and BACE1 under serum deprived conditions, using NP031112, a non-ATP competitive and irreversible GSK-3β inhibitor.^33^ First, we assessed its cytotoxicity after 24 h of treatment and found that concentrations up to 10 μM were non-toxic (Fig. 1B), which were subsequently used for further analyses on SH-SY5Y cells. Western blot (WB) assays were performed to evaluate the protein levels of ATF4, BACE1 and pS9-GSK-3β in the presence of NP031112. As expected, we observed a dose-dependent reduction in ATF4 and BACE1 protein levels, reaching a maximum decrease of over 50% at the highest NP031112 concentration tested (*P* < 0.001) (Fig. 1C). Similar results were obtained with AR·A014418, another GSK-3β inhibitor (Supplementary Fig. 1). Moreover, pS9-GSK-3β levels increased proportionally to the inhibitor concentration, becoming significant (*P* < 0.001) at 10 µM (Fig. 1C). This observation supports the established mechanism where GSK-3β sustains its active state through the inhibitory phosphorylation of I-2, a PP1 inhibitor, consequently allowing PP1 to dephosphorylate inactive pS9-GSK-3β.^34^ Notably, NP031112 appears to interrupt this positive feedback loop by directly inhibiting GSK-3β, offering a potential explanation for the observed elevation in pS9-GSK-3β. This phenomenon is cohesive with the PI3K/Akt-independent increase in Ser9 phosphorylation observed in cells treated with LiCl.^34^ However, at the lowest inhibitor concentrations tested (1 and 5 µM), the difference was not statistically significant, likely due to being insufficient to prevent PP1 activity (Fig. 1C).

**Fig. 1.**
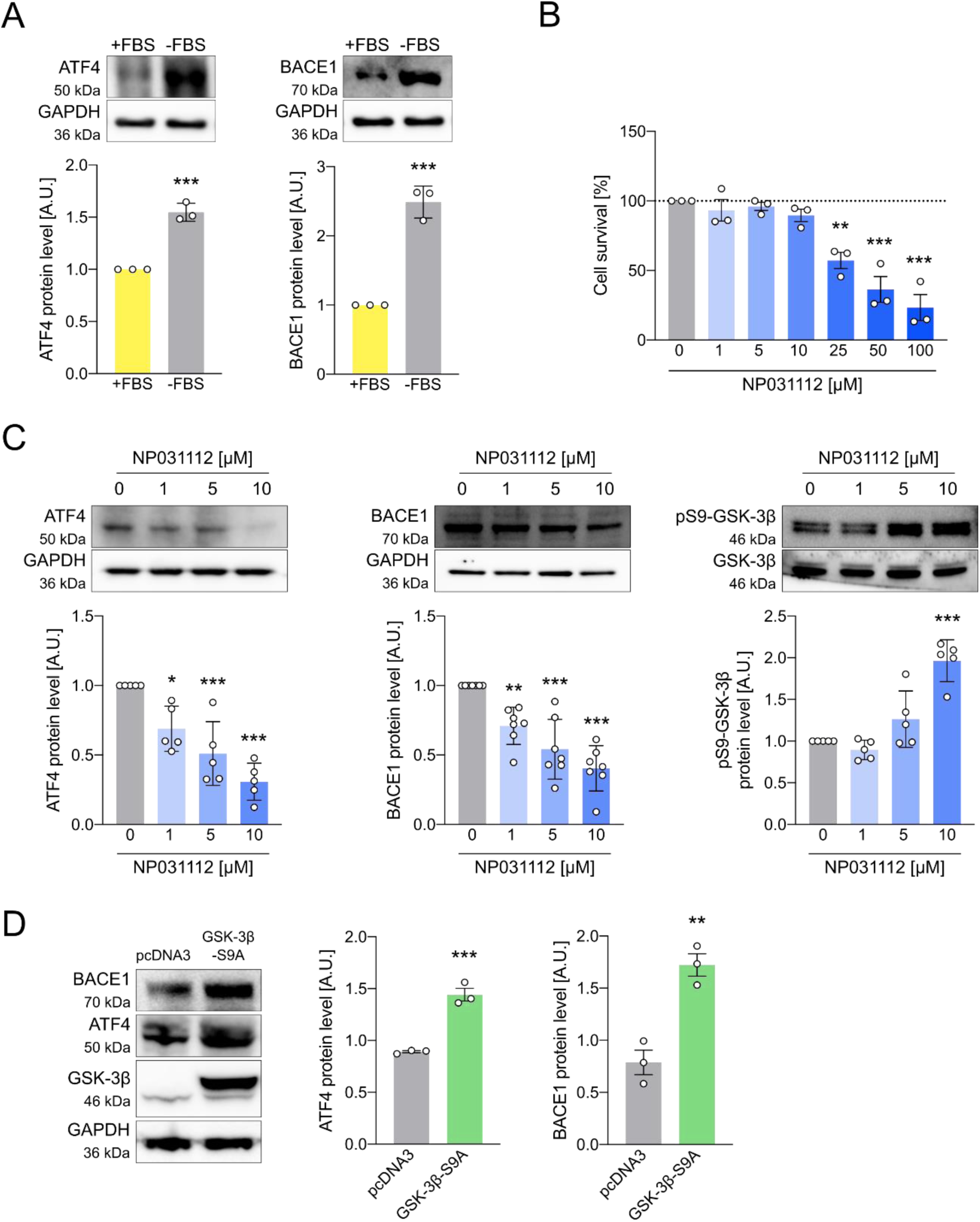
ISR increases ATF4 and BACE1 protein expression through GSK-3β. (**A**) Human neuroblastoma cells (SH-SY5Y cells) were grown for 24 h in the presence/absence of FBS. ATF4 and BACE1 expression was assayed by WB. Data are the mean ± SD of three independent experiments. *** *P* < 0.001 vs. control by Student’s t-test. (**B**) SH-SY5Y cells were treated with increasing concentrations of the GSK-3β inhibitor NP031112 in FBS-free medium for 24 h and survival was assayed by MTT reduction. Data are the mean ± SD of three independent experiments. ** *P* < 0.01 and *** *P* < 0.001 vs. control by one-way ANOVA plus Dunnett as post hoc test. (**C**) SH-SY5Y cells were treated with increasing concentrations of NP031112 in FBS-free medium for 24 h and the expression of ATF4, BACE1 and pS9-GSK-3β were assayed by WB. Data are the mean ± SD of five-seven independent experiments, respectively. * *P* < 0.05, ** *P* < 0.01 and *** *P* < 0.001 vs. control by one-way ANOVA plus Dunnett as post hoc test. (**D**) HEK-293 cells were transfected with empty (pcDNA3) or GSK-3β-S9A-encoding plasmids and BACE1 and ATF4 protein levels were analysed by WB after 24 h. Data are the mean ± SD of four and three independent experiments, respectively. * *P* < 0.05 and *** *P* < 0.001 vs. control by Student’s t-test.

To confirm the specific effect of GSK-3β on the results obtained in Fig. 1A and 1C, we transfected HEK-293 cells with a constitutively active non-phosphorylatable GSK-3β mutant, GSK-3β-S9A, encoding vector. In agreement with prior results, ATF4 and BACE1 levels were significantly increased (*P* < 0.05 and *P* < 0.001, respectively) in cells overexpressing the mutant kinase and kept in FBS-containing media (Fig. 1D). Cohesively, overexpressing the wild-type enzyme also resulted in higher levels of ATF4 and BACE1 (*P* < 0.001 and *P* < 0.01, respectively) (Supplementary Fig. 2); however, these levels were lower than those obtained with GSK-3β-S9A, as expected.

### GSK-3β modulates BACE1 translation and promotes eIF2Bε phosphorylation

To determine whether the increase in BACE1 expression is due to transcriptional regulation, we used 5,6-dichloro-1-beta-D-ribofuranosylbenzimidazole (DRB), a general transcription inhibitor that blocks activating phosphorylation of the RNA polymerase II by inhibiting CDK9, and, therefore, transcript elongation.^35^ Given the high toxicity associated with transcriptional inhibition,^36^ and since DRB acts rapidly (within 5 min)^37^ and does not induce cell death at 5 h, we shortened the treatment duration from 24 to 5 h, also because our results show that GSK-β inhibition decreases BACE1 at 3-5 h (Supplementary Fig. 3). As expected, WB results revealed that both NP031112 and DRB independently reduce BACE1 levels (*P* < 0.001) (Fig. 2A). Notably, a more pronounced reduction in BACE1 levels was observed when both treatments were combined (*P* < 0.001), suggesting that NP031112 still manages to modulate BACE1 expression even when transcription is blocked (Fig. 2A). This observation supports a transcription-independent effect of GSK-3β on BACE1 expression.

**Fig. 2.**
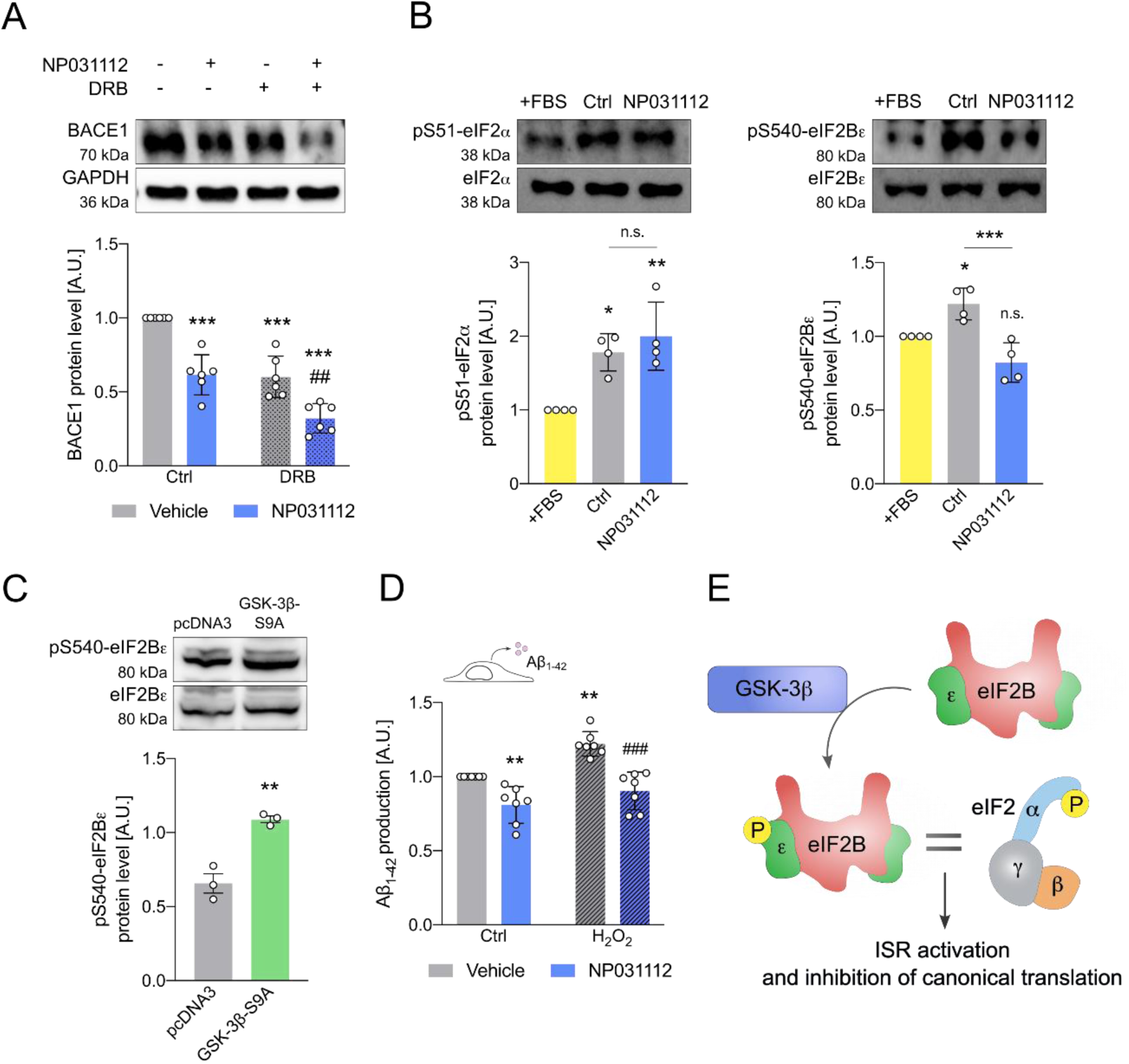
BACE1 up-regulation depends on eIF2Bε phosphorylation. (**A**) SH-SY5Y cells were treated with either 10 μM NP031112, 50 μM DRB, or a combination of both in FBS-free medium for 5 h. The expression of BACE1 was assayed by WB. Data are the mean ± SD of six independent experiments. *** *P* < 0.001 vs. untreated controls and ## *P* < 0.01 vs. DRB plus vehicle by two way-ANOVA plus Tukey as post hoc test. (**B**) SH-SY5Y cells were cultured in FBS-containing medium, serum-starved (Ctrl), or treated with 10 μM NP031112 in FBS-free medium for 24 h and the phosphorylation of eIF2α and eIF2Bε was assayed by WB. Data are the mean ± SD of four independent experiments, respectively. * *P* < 0.05, ** *P* < 0.01, *** *P* < 0.001 and non-significant (n.s.) vs. control by one way-ANOVA plus Dunnett as post hoc test. (**C**) HEK-293 cells were transfected with pcDNA3 and GSK-3β-S9A plasmid and the level of pS540-eIF2Bε was analysed by WB 24 h after transfection. Data are the mean ± SD of three independent experiments. * *P* < 0.05 vs. control by Student’s t-test. (**D**) SH-SY5Y cells were treated with 10 μM NP031112, 10 μM H_2_O_2_, or a combination of both in FBS-free medium for 24 h. Aβ_1-42_ concentration in the supernatant was measured by ELISA. Data are the mean ± SD of seven independent experiments. ** *P* < 0.01 vs. control and ### *P* < 0.001 vs. H_2_O_2_ plus vehicle by two way-ANOVA plus Tukey as post hoc test. (**E**) Schematic representation of the hypothesis: Phosphorylation of the eIF2Bε subunit at Ser540 by GSK-3β mimics the effects of p-eIF2α, due to reduced levels of active eIF2B.

eIF2α is one of the key regulators of the ISR, and its phosphorylation at Serine 51 occurs when one of the four ISR kinases is activated.^13^ Although GSK-3β has not been reported to be an eIF2α kinase, we aimed to determine whether GSK-3β inhibition might influence eIF2α phosphorylation, and therefore explain the typical ISR outcomes we have observed. WB analysis showed a significant increase in eIF2α phosphorylation under serum deprivation (*P* < 0.05). Importantly, GSK-3β inhibition did not revert this, proving that blocking GSK-3β does not affect eIF2α phosphorylation triggered by serum deprivation (Fig. 2B, left panel). Therefore, the increase in pS51-eIF2α observed after serum starvation is independent of GSK-3β overactivation, but rather to GCN2 activation, as GCN2 is the eIF2α kinase that responds to amino acid deprivation.^38^ While no direct link between eIF2α and GSK-3β has been reported, previous studies have shown that GSK-3β phosphorylates eIF2B at Serine 540 of the ε subunit in humans, reducing its activity by approximately 50%.^22,23^ After investigating whether the GSK-3β inhibitor affected eIF2Bε phosphorylation, we found that 10 μM NP031112 significantly reduced pS540-eIF2Bε levels elevated by serum deprivation (*P* < 0.001) (Fig. 2B, right panel). This indicates that the decrease in both ATF4 and BACE1 levels in human neuroblastoma cells observed upon treatment with the GSK-3β inhibitor (Fig. 1) are not caused by a reduction of the canonical eIF2α phosphorylation. Overexpression of the GSK-3β-S9A mutant in HEK-293 cells also increased eIF2Bε phosphorylation (*P* < 0.05), further confirming our findings (Fig. 2C).

Finally, we addressed the pathophysiological relevance of our hypothesis by measuring secreted Aβ_1-42_ (Fig. 2D), the most amyloidogenic Aβ species,^39,40^ in the supernatant of cultured SH-SY5Y cells. To induce Aβ_1-42_ production, we treated human neuroblastoma cells with H_2_O_2_, a well-known GSK-3β activator.^41^ In serum-free conditions, 10 μM NP031112 reduced Aβ_1-42_ levels by approximately 20% (*P* < 0.01). Cells challenged with 10 μM H_2_O_2_ exhibited a 23% increase in Aβ_1-42_ production compared to controls (*P* < 0.01), which was reversed by co-treatment with NP031112 (*P* < 0.001). These findings are consistent with the previous results, showing that less BACE1 implies less Aβ_1-42_ production, and confirms that GSK-3β inhibition contributes to reduced Aβ generation, a result previously observed in mice models.^42,43^

While GSK-3β induces Aβ_1-42_ generation by increasing BACE1 transcription,^27^ indirectly enhancing γ-secretase activity,^44^ and phosphorylating APP and increasing its processing ,^45,46^ no modulation of BACE1 translation by GSK-3β has been established yet. Taken together, our results suggest a model in which GSK-3β modulates eIF2Bε phosphorylation and activates a non-canonical ISR, which at the same time upregulates BACE1 translation, in a situation analogous to that caused by eIF2alpha phosphorylation, which also culminates in a reduction of available eIF2B for protein translation (Fig. 2E). GSK-3β inhibition reduces eIF2Bε phosphorylation, relieving its suppressive effect on global translation initiation, thereby decreasing ATF4 and BACE1 synthesis. This mechanism supports a translational control axis of BACE1 expression and amyloid pathology involving GSK-3β-eIF2Bε-BACE1-Aβ_1-42_.

### An interaction model reveals that eIF2Bε phosphorylation disrupts eIF2 and eIF2B dynamics

To study possible effects on eIF2 and eIF2B dynamics mediated by eIF2Bε phosphorylation, we decided to model their interaction. The structural dynamics of the eIF2-eIF2B interaction are central to translational control via guanine nucleotide exchange mediated by the eIF2B GEF activity. The active structure of eIF2 bound to tRNA (PDB: 3JAP) reveals that GTP is primarily coordinated by the γ subunit, while the β subunit forms a closed conformation over the bound tRNA. The α subunit also participates in stabilizing the tRNA interaction, together forming a ternary complex that is competent for ribosomal recruitment and translation initiation (Supplementary Fig. 4A). Upon GTP hydrolysis, GDP remains tightly bound, likely due to the continued closed conformation of the β subunit, thereby preventing spontaneous nucleotide exchange (Fig. 3A, upper panel).

**Fig. 3.**
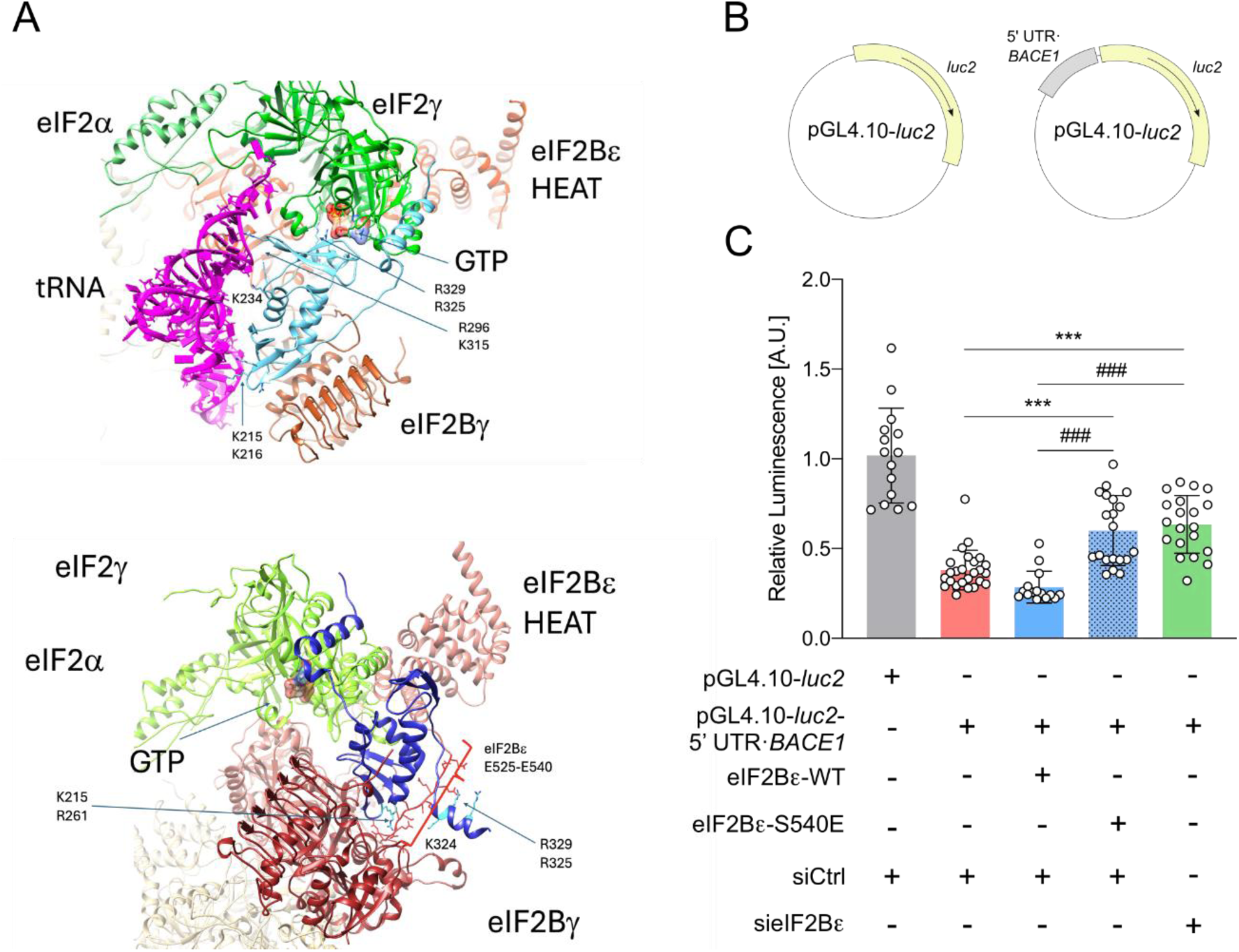
eIF2Bε modelling and mutations support its role on BACE1 translation. (**A**) Ribbon representation of the structural model of eIF2 (subunits α, β, γ) in complex with eIF2B in the closed (up) and open (down) conformations using ChimeraX.^58^ eIF2 subunits α and γ are shown in green; subunit β is shown in cyan in the upper panel and blue in the lower panel. eIF2B subunits γ and ε are shown in dark and light brown, respectively. The remaining eIF2B subunits are shown in beige. The HEAT domain and the GTP molecule are indicated, with the latter displayed in surface representation and coloured by heteroatom. In the upper panel, a tRNA molecule is included to indicate its ability to bind the complex. In the lower panel, the region spanning residues 525-540, enriched in negatively charged residues (including glutamate substitutions for phosphorylated serines), is highlighted in red with side-chain atoms shown. Side-chain atoms of selected positively charged residues (Arg and Lys) are indicated in both panels. (**B**) Schematic representation of the pGL4.10-*luc2* vectors used. (**C**) HEK-293 cells were transfected with vectors pGL4.10-*luc2* or pGL4.10-*luc2*-5’ UTR-*BACE1*; pRL; eIF2Bε-WT or eIF2Bε-S540E; siCtrl or sieIF2Bε. Then, firefly and renilla luciferase activities were analysed with a Dual-Glo luciferase kit and luminescence was recorded with a luminometer. Data are the mean ± SD of 15-25 independent experiments. *** *P* < 0.001 vs. pGL4.10-*luc2*-5’ UTR-*BACE1* or ### *P* < 0.001 vs. eIF2Bε-WT by one-way ANOVA plus Tukey as post hoc test.

The GEF activity of eIF2B is required for GDP release. Structural insights from the eIF2B-eIF2 complex (PDB: 6K71) show that the α subunit of eIF2 adopts an open conformation, while the β subunit is unresolved, suggesting structural flexibility. Superposition of the eIF2-tRNA complex (3JAP) onto the eIF2B-eIF2 complex (6K71) indicates that tRNA could still be accommodated in the complex during nucleotide exchange. The ε and γ subunits of eIF2B mediate the core interaction with eIF2 (Fig. 3A, upper panel).

Using AlphaFold3,^47^ we modelled partial eIF2-eIF2B complexes comprising only the γ and ε subunits of eIF2B, and GDP. Two conformations emerged: (i) a closed β subunit of eIF2 retaining GDP within the active site, with the HEAT domain of the ε subunit distant; and (ii) an open conformation, with the β subunit displaced toward the γ subunit of eIF2B and in proximity to the HEAT domain of eIF2Bε, exposing the GDP-binding site. AlphaFold3 predictions suggest that the interaction favours the open β conformation, facilitating GDP release and subsequent GTP/tRNA binding (Supplementary Fig. 4B).

This supports a model in which eIF2B binds the GDP-bound form of eIF2 and induces opening of the β subunit to enable GDP release. Subsequent GTP loading and tRNA binding promote re-closure of the β subunit, stabilizing the ternary complex. This conformational change is likely mediated by electrostatic interactions, particularly involving positively charged residues on eIF2β (e.g., R261, R296, K215, K234, K259, K315), with R325 and R329 involved in GTP coordination. The α subunit then locks the tRNA in place, and the active ternary complex is released from eIF2B (Supplementary Fig. 4B).

To examine the impact of regulatory phosphorylation, we modelled mutations in the ε subunit of eIF2B that mimic phosphorylation (S525E, S535E and S539E). These mutations induced local disorder in the region connecting the HEAT domain to the core of the ε subunit, as reflected by low pLDDT scores. Despite the disorder, all AlphaFold models predicted an open conformation of the eIF2 β subunit, exposing the GDP-binding site. This conformation appears to result from electrostatic interactions between negatively charged glutamates from eIF2Bε (mimicking the effects of serine phosphorylation) and positively charged residues on eIF2β (notably K215, K259, R261, K324, R325 and R329) (Fig. 3A, lower panel).

Functionally, this phosphorylation mimic mutants disrupts the normal closing of the β subunit, thereby impairing GTP binding and subsequent tRNA capture. The negative charges introduced by phosphorylation compete with the tRNA own phosphate backbone for binding to key basic residues, preventing the formation of a stable ternary complex (Supplementary Fig. 4C). As a result, phosphorylated eIF2 remains in an inactive, GDP-bound state and is unable to initiate translation effectively. These results explain a potential mechanism on how GSK-3β-mediated eIF2Bε phosphorylation achieves ISR activation.

### Inhibition of eIF2Bε enhances BACE1 translation

Next, we studied the effect of eIF2Bε inhibition on BACE1 levels to directly demonstrate a link between eIF2Bε and BACE1 translation and to show the functional relationship between the two. We cloned *BACE1* 5’ UTR into a pGL4.10-*luc2* vector containing the firefly luciferase gene under the control of the SV40 promoter (Fig. 3B). Moreover, to modulate eIF2Bε activity, we used the eIF2Bε-S540E mutant, where the serine at position 540 was replaced by a glutamic acid to structurally mimic the phosphorylation induced by GSK-3β, and a siRNA proven to repress eIF2Bε expression (Supplementary Fig. 5). Compared to the control vector, the insertion of the 5’ UTR reduced the expression of luciferase, as expected (*P* < 0.001) (Fig. 3C). Silencing eIF2Bε, or overexpressing the eIF2Bε-S540E phosphomimetic mutant, rescued luciferase expression in 5’ UTR-expressing cells (*P* < 0.001) (Fig. 3C). These results demonstrate that a reduction of functional eIF2Bε levels induces BACE1 translation, similar to what occurs when eIF2α is phosphorylated.

### Phosphorylated eIF2Bε is increased in Alzheimer’s disease hippocampal samples

To assess whether the molecular alterations observed in culture are relevant for Alzheimer’s disease etiopathogenesis, we analysed post-mortem hippocampal samples from Alzheimer’s disease patients and age-matched non-demented controls. Our study focused on evaluating GSK-3β activation status by analysing its inhibitory phosphorylation at Ser9 (pS9-GSK-3β) and activatory phosphorylation at Tyr216 (pY216-GSK-3β), as well as the expression of downstream targets related to the ISR.

Immunohistofluorescence studies performed on human hippocampal slices revealed a significantly lower pS9-GSK-3β/GSK-3β ratio in Alzheimer’s disease hippocampi compared to controls (*P* < 0.001). In parallel, the pY216-GSK-3β/GSK-3β ratio was found to be significantly higher in Alzheimer’s disease hippocampi (*P* < 0.001), supporting increased GSK-3β activation. These results were confirmed by quantitative analysis of mean fluorescence intensities for both total and phosphorylated forms of GSK-3β (Fig. 4A).

**Fig 4.**
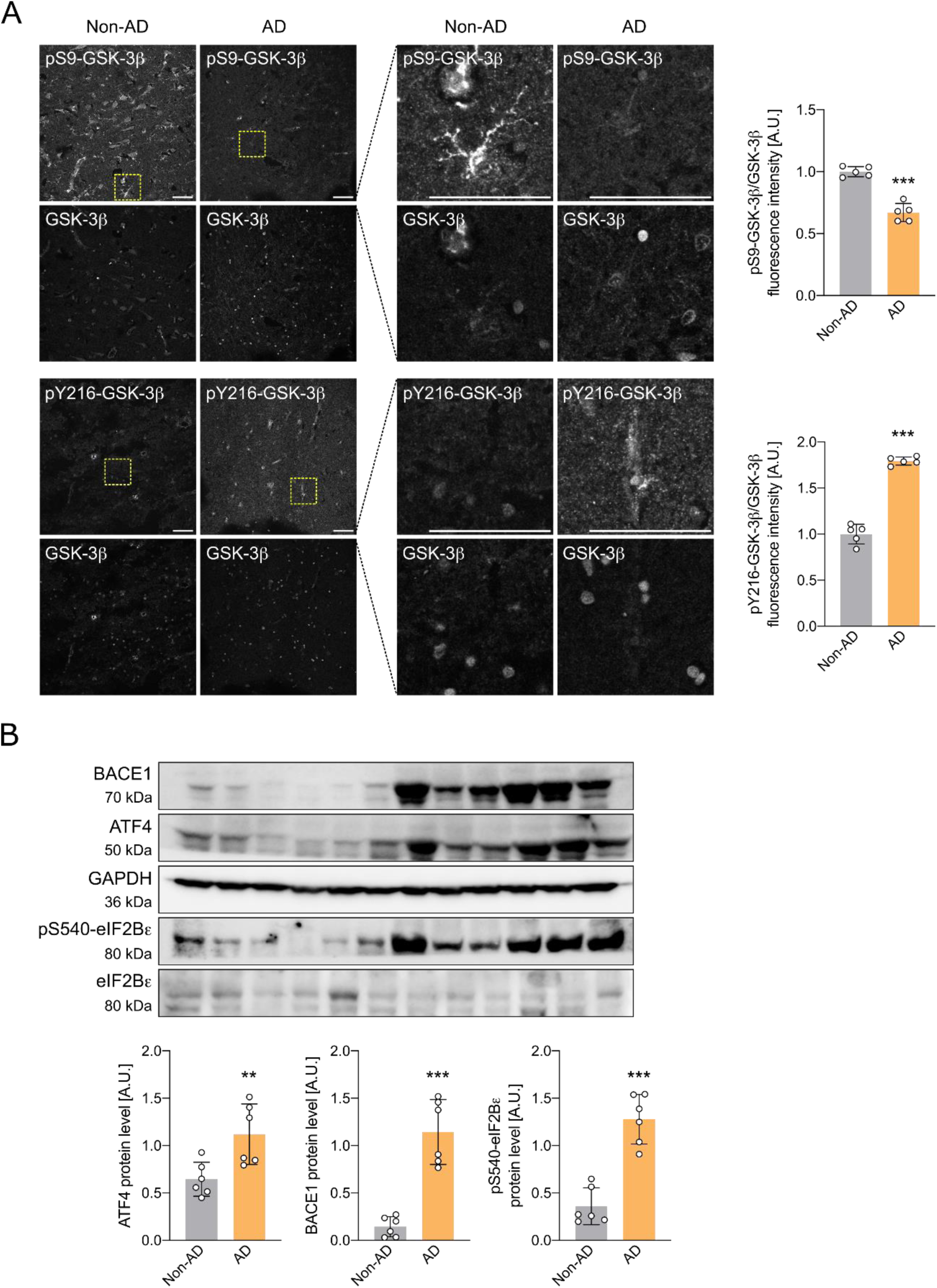
BACE1 and eIF2Bε are increased in the Alzheimer’s disease hippocampal samples. (**A**) Immunohistofluorescence studies of hippocampal slices of Alzheimer’s disease and non-demented individuals were used to investigate GSK-3β activity. The panels on the right show enlarged views of the regions within the yellow-outlined squares in the images on the left panels. Scale bars represent 50 µm in both left and right panels. pS9-GSK-3β and pY216-GSK-3β fluorescence intensities were measured and relativized to the total GSK-3β to create ratios. Data are the mean ± SD of five independent individuals per group. *** *P* < 0.001 vs. control by Student’s t-test. Images were obtained using a Leica SP8 confocal microscope and analysed with ImageJ. (**B**) Hippocampal tissue of Alzheimer’s disease and non-demented individuals was homogenized and levels of ATF4, BACE1 and pS540-eIF2Bε were analysed by WB. Data are the mean ± SD of six independent individuals per group. ** *P* < 0.01 and *** *P* < 0.001 vs. control by Student’s t-test.

To explore the downstream consequences of increased GSK-3β activity, we next analysed the protein levels of key ISR-related components by WB. Hippocampal lysates from Alzheimer’s disease and non-demented individuals were used to quantify pS540-eIF2Bε, ATF4 and BACE1. Consistent with enhanced ISR signalling, we observed a significant increase in ATF4, BACE1 and pS540-eIF2Bε levels in Alzheimer’s disease hippocampi compared to controls (*P* < 0.01, *P* < 0.001 and *P* < 0.001, respectively) (Fig. 4B). These data highlight the relevance of the phosphorylation of eIF2Bε and its potential role in promoting pathological translation of stress-related proteins in Alzheimer’s disease.

Together, these findings provide strong evidence that dysregulation of GSK-3β activity and the ISR axis, particularly via eIF2Bε phosphorylation, contributes to the molecular pathology in Alzheimer’s disease.

## Discussion

Our study uncovers a previously unrecognized role of the ISR through phosphorylation of the eIF2Bε subunit, establishing a novel translational control mechanism that links cellular stress with amyloidogenic processing. Crucially, this pathway converges on BACE1, the rate-limiting enzyme for Aβ generation, placing it at the centre of Alzheimer’s disease aetiology and progression.

We demonstrate that GSK-3β activity increases under stress-like conditions and is both necessary and sufficient to drive a sustained ISR-like state characterized by elevated ATF4 and BACE1. Importantly, this regulation occurs independently of eIF2α phosphorylation, a canonical ISR hallmark, and instead involves direct phosphorylation of eIF2Bε at Ser540, a modification that selectively impairs global translation while favouring translation of stress-responsive mRNAs, such as ATF4 and BACE1. This alternative mechanism mimics the downstream effects of ISR activation but bypasses traditional upstream sensors, suggesting that chronic GSK-3β activation can keep the cell in a “maladaptive translational state”.

The pathological relevance of this mechanism is underscored by our findings in human Alzheimer’s disease hippocampi, where we observed increased GSK-3β activity alongside elevated p-eIF2Bε, ATF4 and BACE1 levels in the hippocampus of Alzheimer’s disease patients compared to controls. These data support the notion that chronic activation of GSK-3β in the aging or diseased brain perpetuates an active ISR state, driving BACE1 expression and consequently enhancing Aβ production, a central event in Alzheimer’s disease pathogenesis.^48^

This mechanism has several implications for our understanding of Alzheimer’s disease. First, it expands the role of GSK-3β beyond insulin resistance, Wnt signalling and tau phosphorylation,^25^ establishing it as a direct modulator of the translational machinery in neurons. Second, it identifies eIF2Bε phosphorylation as a critical node linking stress, translation and amyloid pathology.

While some GSK-3β inhibitors have shown promise in preclinical Alzheimer’s disease models,^49^ translating them to clinical approval has proven challenging, as they are tested in Alzheimer’s disease patients at advanced stages of the disease.^50,51^ Our study, however, offers a crucial perspective by highlighting that the GSK-3β effect on eIF2Bε phosphorylation represent a fundamental mechanism directly impacting protein translation and Alzheimer’s disease aetiology. Specifically, we demonstrate that eIF2Bε phosphorylation leads to increased BACE1 levels and, consequently, enhanced amyloid production. These findings open a promising path for further studies on the regulation of eIF2B activity.

While traditionally viewed as a downstream target inhibited by phosphorylated eIF2α during stress, our research demonstrates that eIF2B itself can act as a trigger of ISR-like responses when inhibited by GSK-3β-mediated phosphorylation. This occurs because the reduced availability of functional eIF2B alters translation dynamics, slowing ribosomal activity in a manner similar to the effects of eIF2α phosphorylation. These results are consistent with recently published evidence that shows that deletion of eIF2B alone promotes the activation of an alternative ISR, inducing a metabolic reprogramming to maintain energy homeostasis upon eIF2B attenuation.^52^ These discoveries shift the perspective on eIF2B from being merely a passive target to an active component in stress sensing and response. Its relevance is further underscored by the fact that mutations in eIF2B subunits cause Vanishing White Matter (VWM) disease, a leukodystrophy marked by white matter degeneration and high vulnerability to physiological stress, particularly in glial cells like astrocytes, which significantly shortens the life expectancy of patients, who have severe neurological symptoms.^53–55^ These mutations often disrupt eIF2B guanine nucleotide exchange activity,^56^ which resembles the phosphorylation effect we observed in our interaction model. On the other hand, the discovery of molecules such as ISRIB, which stabilizes eIF2B and restores its function even under stress,^57^ has opened up new therapeutic possibilities not only for neurodegenerative diseases, but also for disorders directly linked to eIF2B dysfunction.

## Conclusions

Our findings suggest that a decline in functional eIF2Bε levels triggers ISR-associated protein translation, paralleling the mechanism activated by eIF2α phosphorylation. It establishes a novel GSK3β-eIF2Bε-BACE1 axis as a mechanistic link between stress response dysregulation of translation and Alzheimer’s disease pathology. By elucidating how ISR reprograms translation in neurons to favour amyloidogenic pathways, we reveal a central pathogenic mechanism in Alzheimer’s disease aetiology.

## List of abbreviations

Ab: antibody
Aβ: amyloid β-peptide
AD: Alzheimer’s disease
Akt: Protein Kinase B
APP: Amyloid Precursor Protein
ATF4: Activating Transcription Factor 4
A.U.: arbitrary unit
BACE1: β-site APP-cleaving enzyme 1
BSA: Bovine serum albumin
CDK9: Cyclin-Dependent Kinase 9
DRB: 5,6-dichloro-1-β-D-ribofuranosylbenzimidazole
ELISA: Enzyme-Linked Immunosorbent Assay
eIF2α: Eukaryotic Initiation Factor 2 alpha
eIF2B: Eukaryotic Initiation Factor 2B
eIF2Bε: Eukaryotic Initiation Factor 2B epsilon subunit
FBS: foetal bovine serum
GCN2: General Control Nonderepressible 2
GDP: guanosine diphosphate
GEF: Guanine Nucleotide Exchange Factor
GSK-3β: Glycogen Synthase Kinase 3 beta
GTP: guanosine triphosphate
HEAT: (Huntingtin Elongation Factor 3, PR65/A, TOR) domain
HEK-293: Human embryonic kidney 293 cell line
I-2: Inhibitor-2 (of PP1)
ISR: Integrated Stress Response
ISRIB: Integrated Stress Response Inhibitor
MTT: 3-(4,5-dimethylthiazol-2-yl)-2,5-diphenyltetrazolium bromide
n.s.: non-significant
PBS: phosphate-buffered saline
PI3K: Phosphoinositide 3-Kinase
pLDDT: predicted Local Distance Difference Test
PP1: Protein Phosphatase 1
RT: room temperature
SH-SY5Y: Human neuroblastoma cell line
SDS: sodium dodecyl sulphate
tRNA: transfer ribonucleic acid
uORF: Upstream Open Reading Frame
UTR: untranslated region
VWM: Vanishing white matter disease
WB: western blot.

## Supplementary material

Supplementary material is available at *XXXX* online.

## Author contributions

HFU, AA, BO and FJM designed the experiments. HFU, CPC, OB, CPP, ST and BO carried out the experiments. HFU and FJM analyzed and interpreted the data and wrote the manuscript. BO and FJM provided the fundings for the work. All authors reviewed the manuscript.

## Data availability

The data that support the findings of this study are available on reasonable request from the corresponding author.

## SUPPLEMENTARY MATERIALS

**Supplementary Fig. 1.**
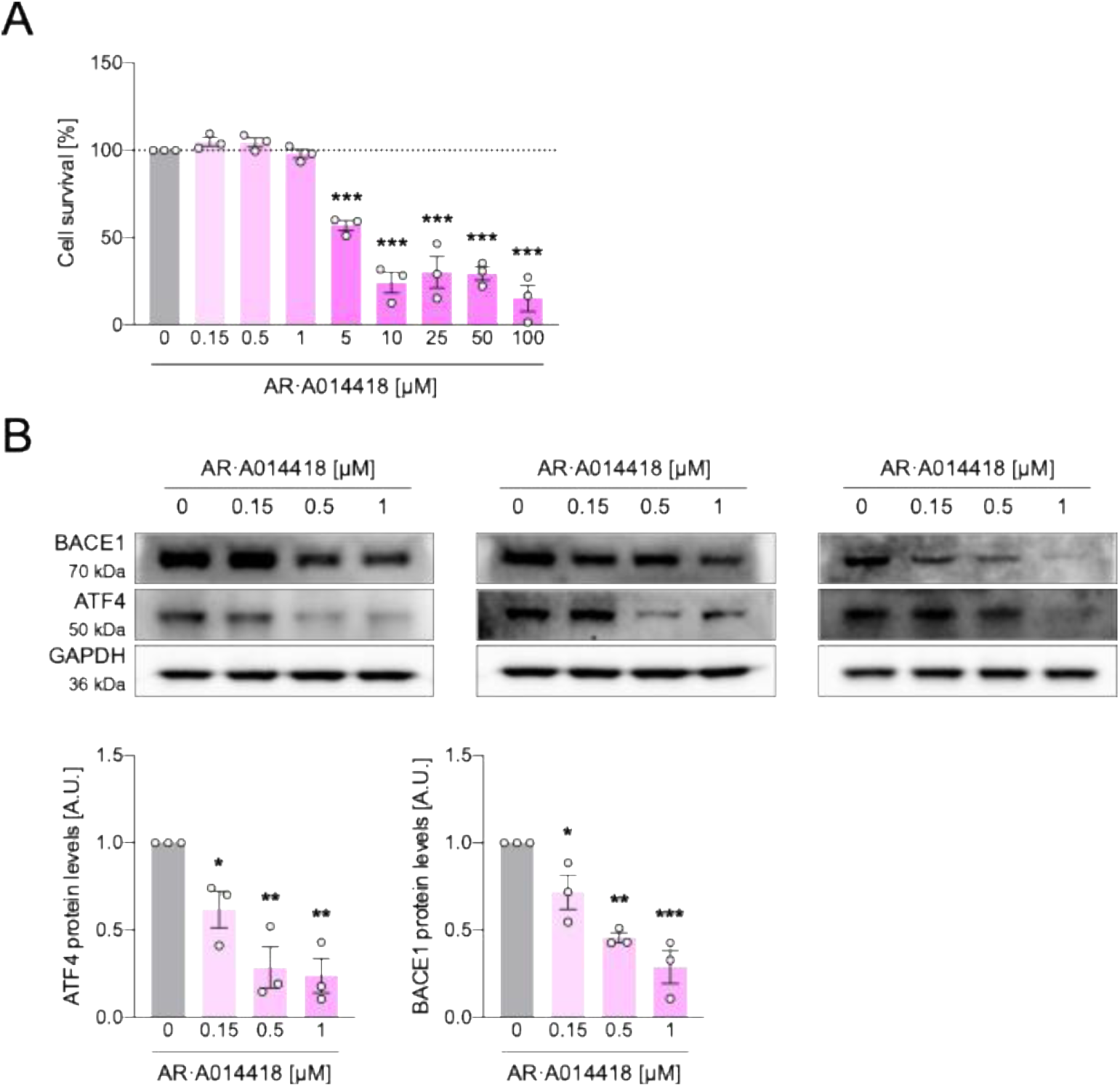
AR·A014418 reduces ATF4 and BACE1 levels. (**A**) SH-SY5Y cells were treated for 24 h with increasing concentrations of AR·A014418. Then, a MTT assay was performed to analyse cell survival. Data are the mean ± SEM of 3 independent experiments, respectively. *** *P* < 0.001 vs. control by one-way ANOVA plus Dunnett as post hoc test. (**B**) SH-SY5Y cells were treated for 24 h with increasing subtoxic concentrations of AR·A014418. Then, WB were performed to analyse ATF4 and BACE1 protein levels. Data are the mean ± SEM of 3 independent experiments, respectively. *** *P* < 0.001, ** *P* < 0.01 and * *P* < 0.05 vs control by one-way ANOVA plus Dunnett as post hoc test.

**Supplementary Fig. 2.**
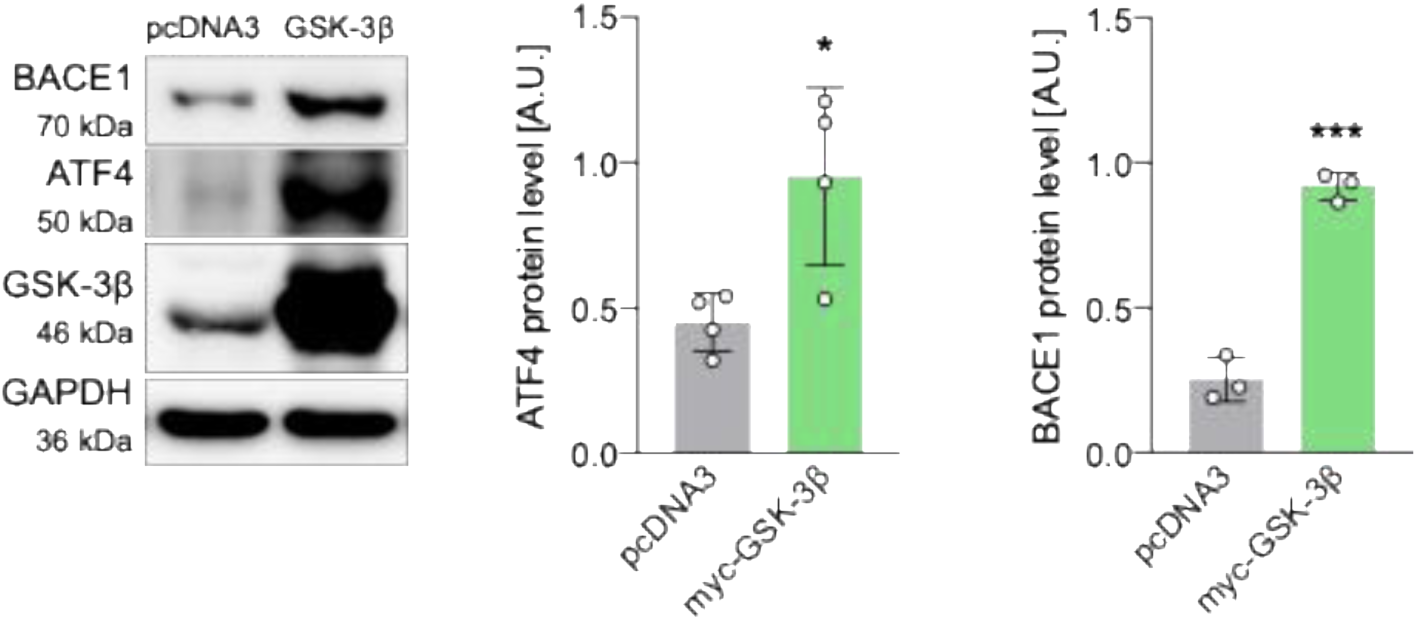
Overexpression of GSK-3β-WT upregulates stress-responsive signalling. HEK-293 cells were transfected with empty (pcDNA3) or GSK-3β-encoding plasmids and BACE1 and ATF4 protein levels were analysed by WB after 24 h. Data are the mean ± SD of four and three independent experiments, respectively. * *P* < 0.05 and *** *P* < 0.001 vs. control by Student’s t-test.

**Supplementary Fig. 3.**
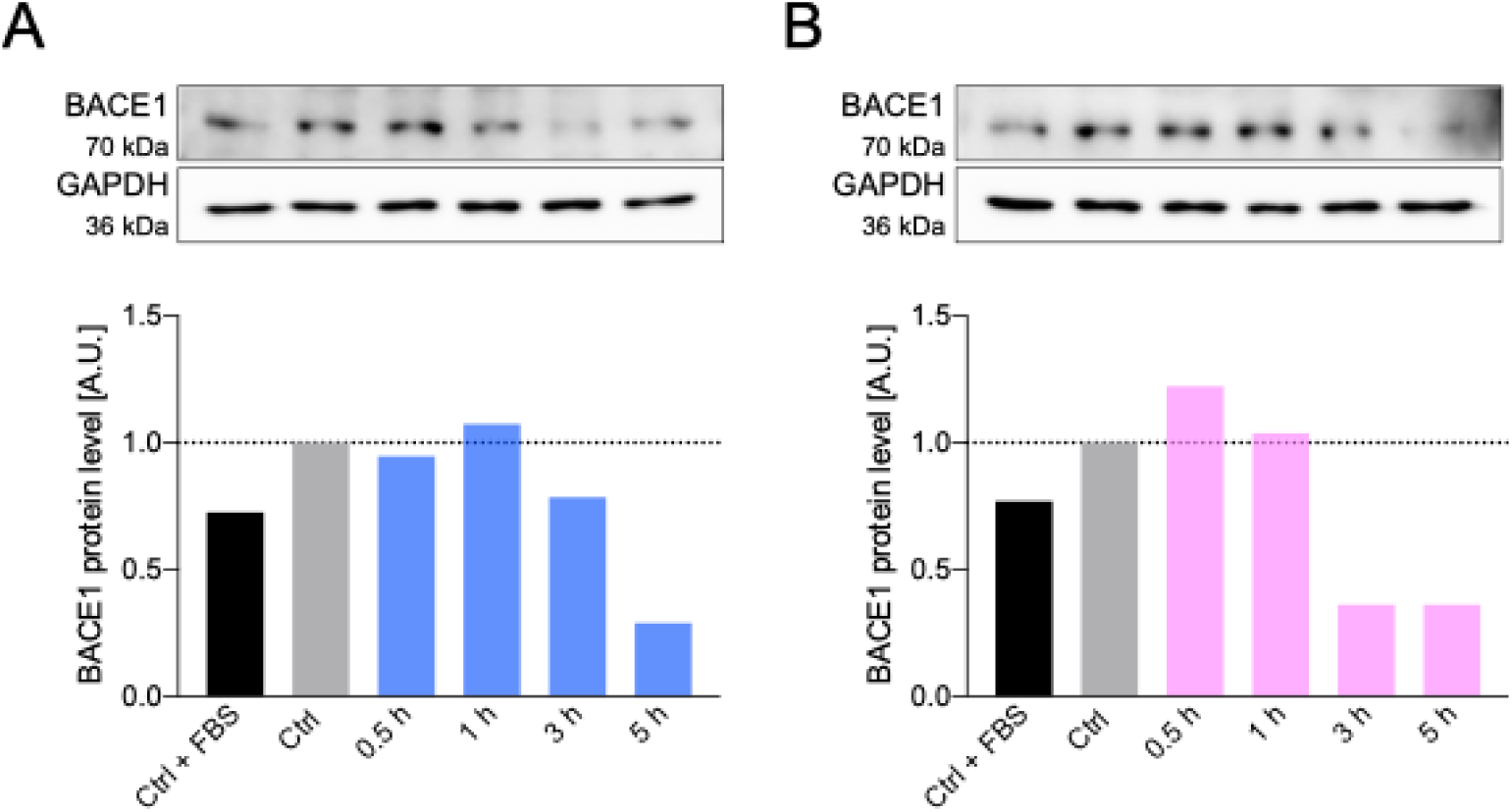
NP031112 and AR·A014418 reduce ATF4 and BACE1 levels. SH-SY5Y cells were treated for 0 (Ctrl + FBS and Ctrl), 0.5, 1, 3 or 5 h with 10 µM NP031112 (**A**) or 1 µM AR·A014418 (**B**). Then, a WB was performed to analyse BACE1 protein levels, and the bands were quantified.

**Supplementary Fig. 4.**
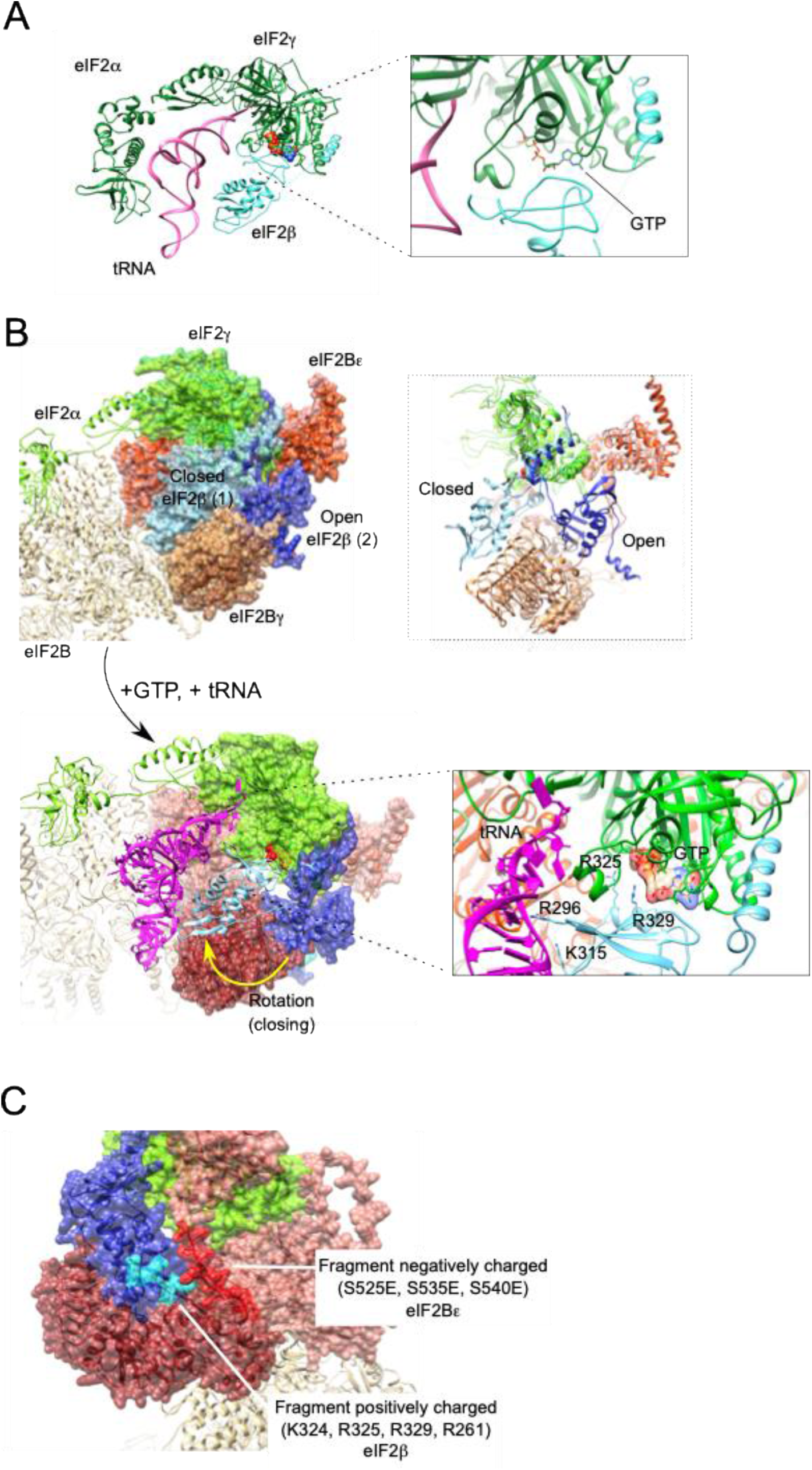
Phosphorylation of eIF2Bε disrupts proper formation of the translation ternary complex. (**A**) Ribbon representation of the structural model of eIF2 (subunits α, β, γ) in complex with a GTP molecule and a tRNA (in pink) using ChimeraX. eIF2 subunits α and γ are shown in green; subunit β is shown in cyan in the upper panel and blue in the lower panel. (**B**) Binding of eIF2B to eIF2 predicts two possible β subunit conformations: 1) a closed conformation shown in cyan; and 2) an open conformation shown in dark blue. The electrostatic interaction between the positively charged residues R296, K315, R325, and R329 on eIF2β and the GTP and tRNA molecules is depicted. eIF2B γ subunit is shown in brown/maroon and ε subunit is shown in red/translucent red. The rest of eIF2B subunits are partially showed in beige. (**C**) Representation of the interaction between eIF2 and eIF2B containing the phosphomimetic glutamate residues in eIF2Bε. eIF2β is represented in dark blue, with positively charged fragment highlighted in cyan; eIF2Bε subunit is shown in translucent red, with negatively charged phosphomimetic residues highlighted in red.

**Supplementary Fig. 5.**
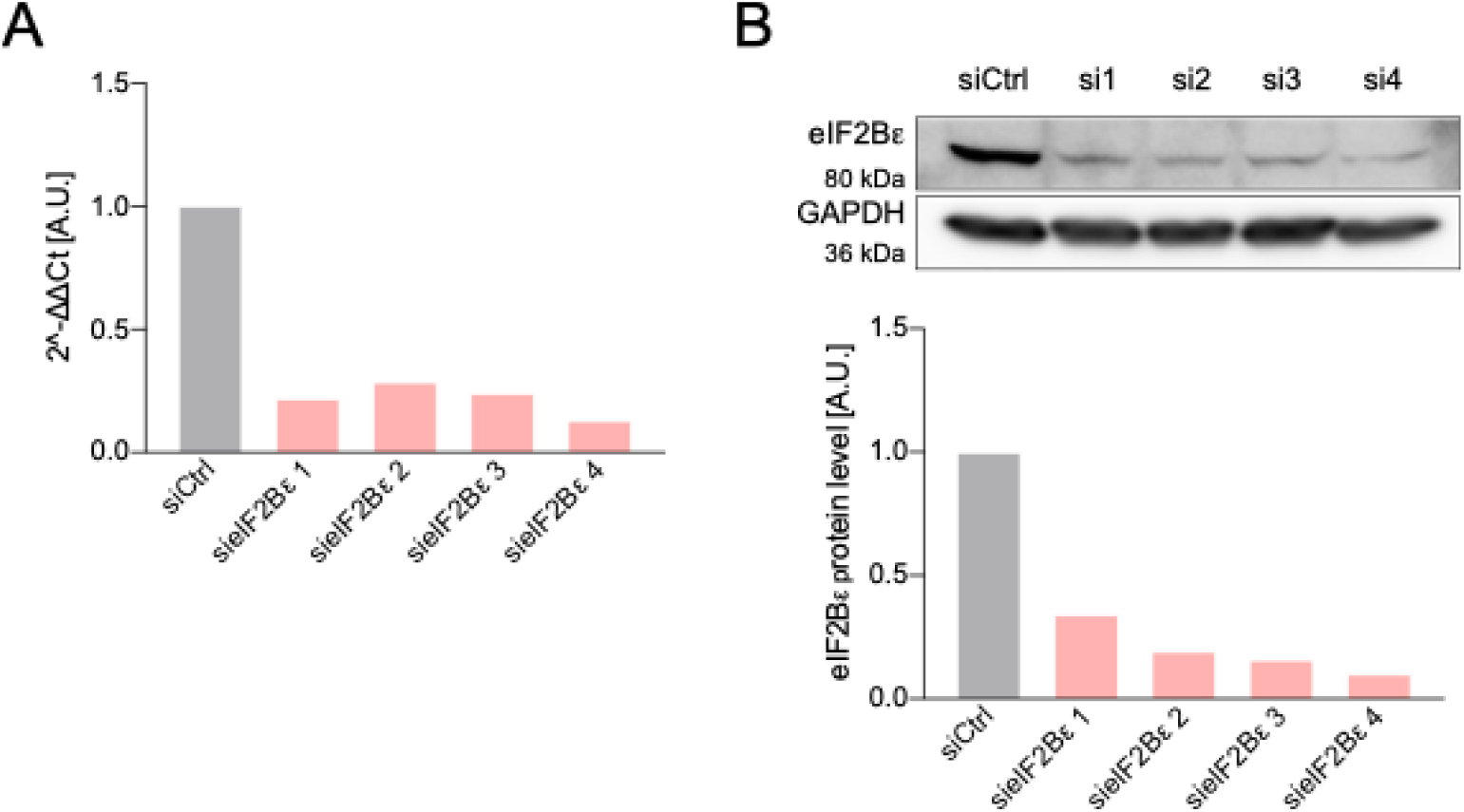
Silencing eIF2Bε reduces its mRNA and protein levels. (**A**) HEK-293 cells were transfected for 24 h with four different siRNAs targeting different sequences of the *EIF2B5* mRNA. Then, a RT-qPCR was performed to study levels of *EIF2B5* mRNA. (**B**) A western blot was also performed to study the protein levels of eIF2Bε after silencing.

